# Functional connectivity of the developing mouse cortex

**DOI:** 10.1101/2021.04.04.438377

**Authors:** Rachel M. Rahn, Lindsey M. Brier, Annie R. Bice, Matthew D. Reisman, Joseph D. Dougherty, Joseph P. Culver

## Abstract

Cross-sectional studies have established a variety of structural, synaptic and cell physiological changes corresponding to key critical periods in cortical development. However, the emergence of functional connectivity (FC) in development has not been fully characterized, and hemodynamic-based measures are vulnerable to any neurovascular coupling changes occurring in parallel. We therefore used optical fluorescence imaging to trace longitudinal calcium FC in the awake, resting-state mouse cortex in the same mice at 5 developmental time points beginning at postnatal day 15 (P15) and ending in early adulthood (P60), resulting in over 500 imaging epochs with both calcium and hemodynamics available as a resource for the field. Proof-of-principle analyses revealed that calcium FC displayed coherent functional maps as early as P15, and FC significantly varied in connections between many regions across development, with the developmental trajectory’s shape specific to the functional region. This longitudinal developmental calcium FC dataset provides an essential resource for further algorithm development and studies of healthy development and neurodevelopmental disorders.

## Introduction

Functional connectivity (FC) imaging is a minimally invasive technique by which circuit and network function can be observed *in vivo* in the human and animal brain. While structural connectivity characterizes intra- and interregional physical connections, FC investigates the functional relationship between neural activity in different brain regions, and has been used extensively in clinical populations as well as in animal models to probe healthy function and disease-related dysfunction. While FC imaging is a powerful tool that provides insight into how the brain functions under a wide variety of conditions, the development of the functional connectome between birth and adulthood is not well-characterized.

Largely cross-sectional FC studies have previously provided insight into health and disease at discrete time points, displaying some sensitivity to development-specific change. The strengthening and reshaping of multiple functional networks, including those involving motor and cingulate regions, has been previously documented across several postnatal weeks in preterm infants and across several months in term human infants (Damaraju et al., 2014; Fransson et al., 2007, 2011; Lin et al., 2008; Smyser et al., 2010, 2011). The strength of functional connections during postnatal development have also been shown to change in both global and region-specific manners, as the ratio of distal to local connection strength has been reported to increase across time in rodents (Ma et al., 2018), and functional lateralization has been documented during language and visuospatial development in humans (Groen et al., 2012). FC has also been found to dynamically change in the visual cortex during the rodent visual critical period and therefore be reflective of well-established cellular-level and functional changes (Kraft et al., 2017). FC measures during early development have also been demonstrated to have a predictive value for the future diagnosis of autism (Emerson et al., 2017), suggesting that FC has strong promise as a biomarker of neurodevelopmental disease. However, although human connectome studies are collecting longitudinal datasets, there currently is a lack of complete longitudinal childhood FC datasets. Furthermore, the blood oxygen level dependent (BOLD) imaging data available represent an indirect measure of neural activity, making untested assumptions that neurovascular coupling is completely established in the developing mammalian brain.

While the emergence of FC throughout development is not fully known, previously documented changes in the structural connectivity of the developing brain indicate that there are both brain-wide and region-specific factors acting after birth and into adulthood. Synaptogenesis, thalamocortical projection reorganization, myelination and synaptic remodeling all have time courses that differ across cortical region and circuit (Huttenlocher and Dabholkar, 1997; Tau and Peterson, 2010). Synaptogenesis in humans peaks in infancy or early childhood (Huttenlocher and Dabholkar, 1997), although the exact progression of synaptic growth and pruning is highly region-dependent. For example, synaptic density is highest in V1 between 4 and 12 months of age in humans (Huttenlocher, 1990), but highest in prefrontal cortex between 8 months and 2 years (Huttenlocher, 1979), suggesting that functional development might likewise display a peak earlier in visual than frontal regions. In mice, synapse density rapidly increases in all brain regions in the first month before fluctuating in a region-specific manner, and synapse composition similarity between brain regions is highest in the first postnatal week in the mouse and then decreases until three months of age (Cizeron et al., 2020). Meanwhile, the time course of synaptic or collateral projection pruning also varies across brain regions, with pruning occurring in the human sensorimotor cortices shortly after birth but later in association cortices and corpus callosum (Huttenlocher and Dabholkar, 1997; Innocenti and Price, 2005). Brain lateralization also occurs during the postnatal period (Andrew 2002), with laterality in rodent cerebral cortex morphology (Diamond et al., 1983) and changes in experience-dependent functional lateralization and callosal development also seen in mice during the developmental period (Bulman-Fleming et al., 1992). Although longitudinal bilateral FC data during development is not available, it is likely that in at least some regions, an increase in structural lateralization is paralleled by decreased bilateral FC. Myelination changes also are well-documented as occurring throughout the developmental period, peaking during the first year of life in humans but continuing into young adulthood, particularly in certain cortical regions (Fields, 2008; Miller et al., 2012; Snaidero and Simons, 2014). In the mouse, myelination begins after birth, continues during adolescence and is nearly complete at postnatal day 60 in most brain regions (Baumann and Pham-Dinh, 2001), suggesting that myelination could be a driving force behind observable FC changes between late adolescence and adulthood. All of these changes in brain structure during development indicate a FC change during this window is also likely and should have some element of region specificity, as functional and structural connections are well-documented to have a close but not perfectly parallel relationship with each other (Damoiseaux and Greicius, 2009; Straathof et al., 2019). However, it is unclear what aspects of structural connectivity most directly contribute to FC, and if therefore changes in synaptogenesis, synaptic pruning, or myelination might have the greatest influence on developmental trajectories. For example, a peak in FC strength early in childhood and then a steady decline might occur if FC closely parallels synaptic density or pruning trends, while a close relationship between FC and myelination might suggest a steep increase in FC early in development and then more gradual but continuing growth in FC strength into adulthood. The difficulty of collecting longitudinal FC during this period has previously prevented studies addressing these questions.

Mouse models and calcium indicators of neural activity provide an excellent opportunity to directly map FC development in the same individuals across time and make new discoveries regarding the brain’s functional development. While the timespan required to collect longitudinal data in humans presents challenges, mice have a comparatively rapid maturation to adulthood as well as well-controlled genetic and environmental factors that reduce variability not attributable to age. Furthermore, mice allow for a more direct measure of neural activity, rather than the use of blood flow as a proxy: Previous research indicates that there may be an inversion in neurovascular coupling and therefore the direction of the hemodynamic response within the first few weeks of life in rodents (Kozberg et al., 2016), making BOLD- and other hemodynamic technique-based studies tracing FC across this period potentially not representative of true neuronal activity change. Previous BOLD-based studies of FC development indicating a strengthening of connections even in infancy may therefore only reflect vascular and not neural changes during this period. Use of a genetically encoded calcium indicator (GECI) as a more direct measure of neuronal activity, however, allows this limitation to be circumvented and the activity of the neural circuits within the brain and their FC to be more directly measured. We therefore collected longitudinal calcium fluorescence FC data using an optical fluorescence imaging system and *Thy1*-GCaMP6 mouse model to directly measure the activity of excitatory pyramidal neurons in the cortex of awake mice in the resting alert state at five developmental time points between postnatal day 15 (P15) and P60 (Figure 1A-C). By characterizing individuals’ neural activity across time, we aimed to establish if early FC development displays similar change to infant FC when calcium neural activity and not BOLD measures are taken. Globally, we hypothesized that a decrease in bilateral FC would parallel the increased lateralization seen in structural connectivity studies, and that overall, FC measures would display region-specific characteristics such as an earlier peak in visual than frontal cortices. Further, we sought to determine if a steady increase after sharp early growth occurs in FC paralleling myelination, or instead if FC has an early peak followed by a decline paralleling synaptic density and pruning time courses. Finally, we examined the data in an unbiased manner to discover additional patterns, as our unique longitudinal functional dataset enables novel discoveries regarding the principles of the development of FC. We therefore have produced a developmental cortical FC dataset representing a total of 510 imaging epochs longitudinally collected from 17 mice at 5 time points.

**Figure 1:**
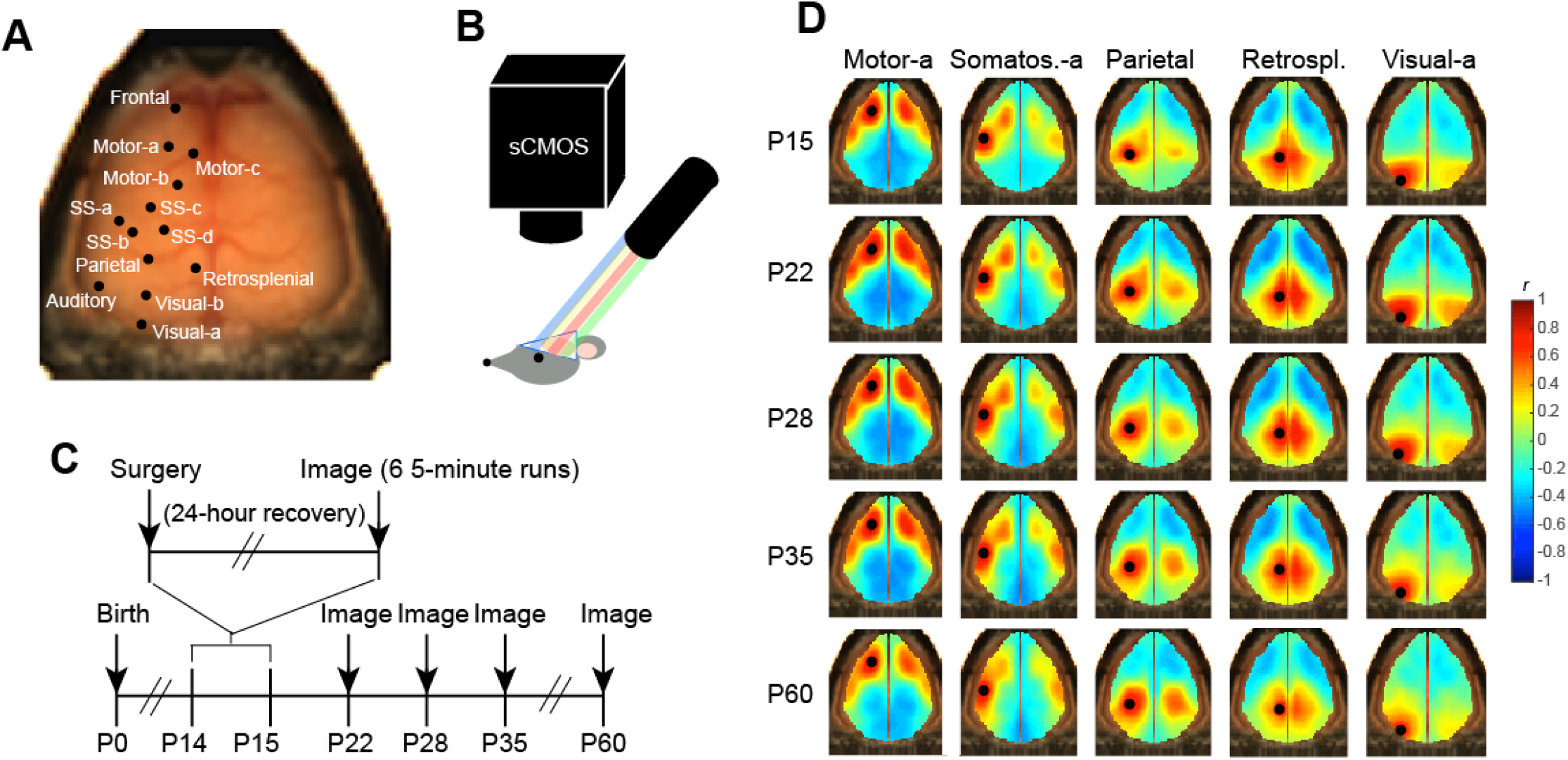
Calcium fluorescence imaging of dorsal cortex provides longitudinal information regarding functional connectivity patterns in mice P15-P60. A) Cortical seed map of the left hemisphere, representing 13 of 26 total cortical seeds. The remaining 13 seeds are mirrored onto the contralateral cortex. B) Schematic of mouse head and imaging system. C) Timeline of experiment, with optical window implantation at P14 followed by longitudinal imaging timepoints at P15, P22, P28, P35, and P60. D) Mean functional connectivity maps in the same cohort across time in multiple sensorimotor and association cortical regions (n=17, 0.4-4.0 Hz).

## Results

### Neuronal activity-dependent FC is established as early as postnatal day 15

Previous optical FC studies have focused primarily on the adult or juvenile stages (Kraft et al., 2017; White et al., 2011; Wright et al., 2017) and rodent fMRI research has used the blood oxygenation level-dependent signal as a correlate of the neuronal signal when examining the development of FC architecture (Ma et al., 2018). Calcium fluorescence recorded by the use of genetically encoded calcium indicators more closely reflects neuronal activity than hemodynamic measures, as neurovascular coupling has been shown to vary according to disease state (Tarantini et al., 2017), but the presence of calcium FC networks as early as P15 has not previously been established and characterized in detail. Therefore, we first examined whether the developmental time points before early adulthood exhibited data quality and general connectivity patterns comparable to adults. Seed maps at P15 already displayed high homotopic contralateral calcium connectivity with the center of these highly-correlated contralateral regions typically having Pearson *r* greater than 0.5, and more distal and functionally unrelated regions having no correlation or an anticorrelation to the seed (Figure 1D, 1st row). Calcium functional connectivity therefore appears to be present and displays patterns typical of adult functional networks as early as P15.

### Homotopic contralateral functional connectivity changes across development in multiple regions

Bilateral functional connections typically mirror the patterns seen in callosal connections, with particularly strong connectivity between homotopic contralateral regions that share functional domains. The corpus callosum displays structural development reflective of phenomena such as synaptogenesis, exuberant projection pruning and myelination, and structural connectivity studies have suggested that connection development is region-specific. We therefore hypothesized that we would also see strong connections between homotopic seeds in left and right hemispheres that would strengthen and then decline, similar to synaptic density and pruning trajectories, with variation dependent on the specific region’s function and related critical period.

We indeed found that the FC developmental trajectories were highly dependent upon the region examined. Bilateral connectivity between the motor seeds displayed an increase between P15 and P22, then a subsequent decrease across the remaining time points, similar to the retrosplenial seeds’ FC trajectory (Figure 2). The visual seeds displayed another type of curved trajectory, as homotopic contralateral connectivity in the visual cortex increased between P15 and 22, then declined to P35 and remained steady between P35 and P60, displaying significant change across time (rmANOVA *p*=4.34e-04(*FDR), pairwise comparisons P15-P22 *p*=2.99e-06*, P22-28 *p*=.0575, P28-P35 *p*=.00752*, P35-P60 *p*=.296) (Figure 2F). Bilateral somatosensory connectivity was largely flat, failing to change significantly across the developmental period. More linear trajectories were observed in frontal regions, where FC largely decreased, and in parietal cortex, which displayed a net increase in FC. The variety in developmental trajectories between homotopic contralateral seed pairs suggests that functional connections develop highly dependent on the region they represent and are not uniform across the cortex. Many regions such as visual, motor, frontal, and retrosplenial cortex, display a peak and then decline similar to the shape of synaptic density and pruning curves.

**Figure 2:**
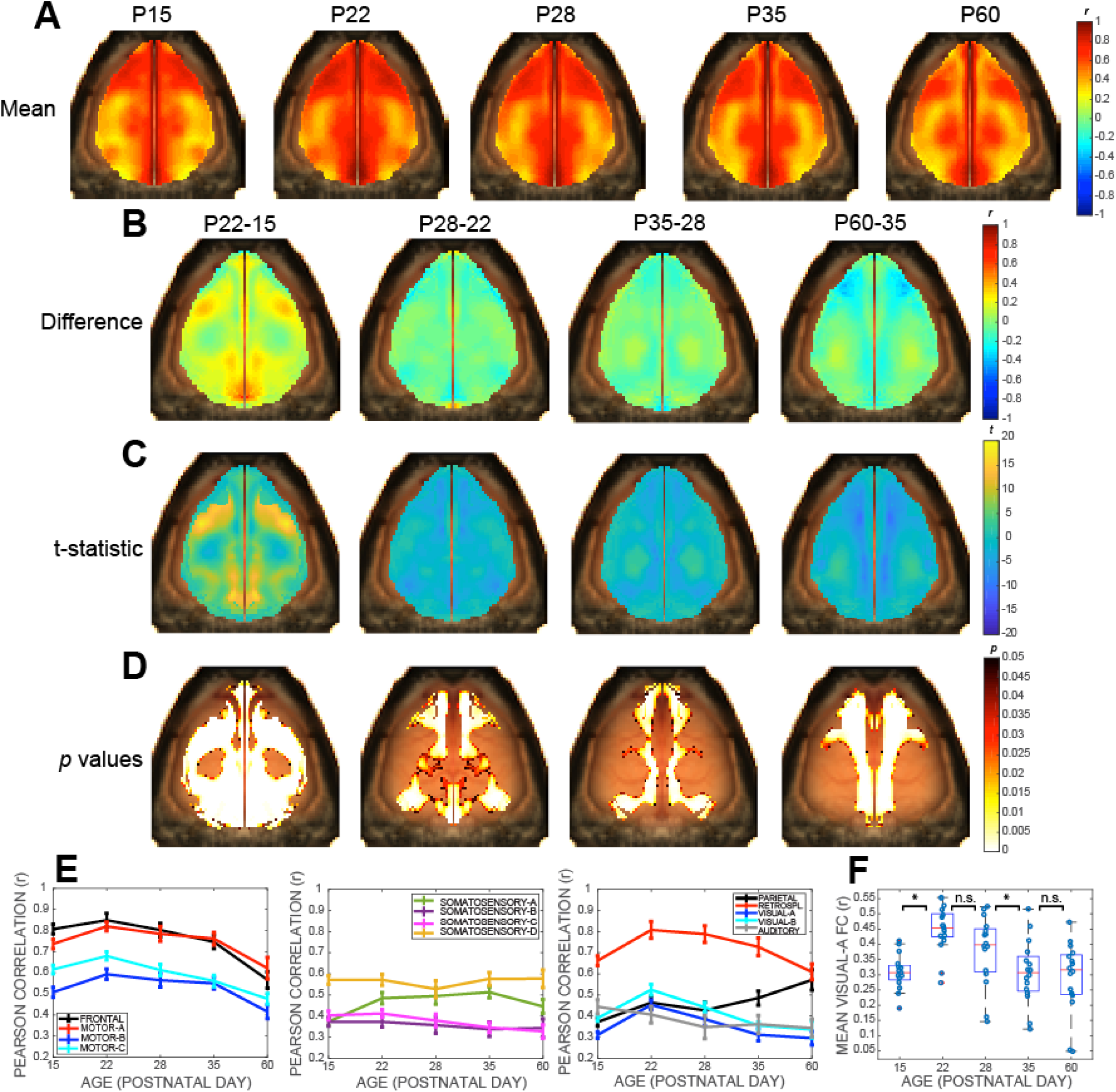
Homotopic contralateral connectivity displays dynamic change across multiple cortical regions during development. A) Mean cortical map of each pixel’s correlation with its homotopic contralateral counterpart across midline at each timepoint. B) Difference between mean bilateral maps at consecutive timepoints, subtracting the earlier time point from the later. C) *t*-statistic maps of paired t-tests between consecutive time points. D) Pixels with significant homotopic contralateral FC changes between consecutive timepoints following a cluster-based multiple comparison correction. E) Mean homotopic contralateral connectivity ±SEM between seed pairs in all 13 cortical regions of interest F) Paired t-test of visual seeds’ connectivity changes between time points (P15-P22 *p*=2.99e-06*, P22-28 *p*=.0575*, P28-P35 *p*=.00752*, P35-P60 *p*=.296) (Pearson *r*, n=17, 0.4-4.0 Hz).

### Both sensorimotor and association cortices display dynamic developmental change in functional connections with other regions

Beyond the 13 homotopic contralateral FC seed pairs, an additional 312 seed-to-seed comparisons within the cortex can be examined to analyze a total set of 325 functional connections and identify regions that are hubs of developmental change. This represents a newly expanded seed set over our prior analyses (Bauer et al., 2014; White et al., 2011) with up to 4 seeds per cortical regions demarcated with lowercase letters (e.g. SS-a, SS-b). Because synaptic pruning has been reported to occur later in association cortex and the corpus callosum than sensorimotor areas, we hypothesized that regions such as retrosplenial and frontal cortices would display more dynamic change than sensorimotor regions. We hypothesized that the sensorimotor regions would have already experienced a significant amount of synaptic pruning before the third postnatal week when we begin sampling, and that bilateral connections would display more change across time because of the later corpus callosum projection pruning. Following a Bonferroni correction for multiple comparisons, we observed that each of the 26 cortical seeds displayed significant change across time for at least 7 of each seed’s 25 functional connections, with the Motor-b, Somatosensory-d and Visual-b seeds generally displaying the highest number of significant connections (Figure 3A,B). A separation between association and sensorimotor regions, as hypothesized, did not widely occur: The aforementioned sensorimotor seeds had significant development in their FC with other regions, and association cortex as represented by parietal, frontal, and retrosplenial seeds also had alterations in FC with other seeds. While somatosensory seeds did not have significant developmental trajectories in their bilateral connectivity across time, somatosensory cortex did significantly change in connections with visual and auditory seeds during the period sampled. Just as synaptic density and pruning display a clear peak in structural literature, a common peak appeared to occur in many functional connections: Dynamic change was especially evident in many connections before and after P22 (Figure 3C-F), with the P15-P22 and P22-P28 periods displaying many significantly altered connections in pairwise comparisons. Overall, functional connections appeared to strengthen between P15 and P22, with a decline in connections between P35 and P60 in the frontal and motor seed sets among others, with every cortical region displaying significant developmental trajectories with both homotopic and non-homotopic cortical seeds.

**Figure 3:**
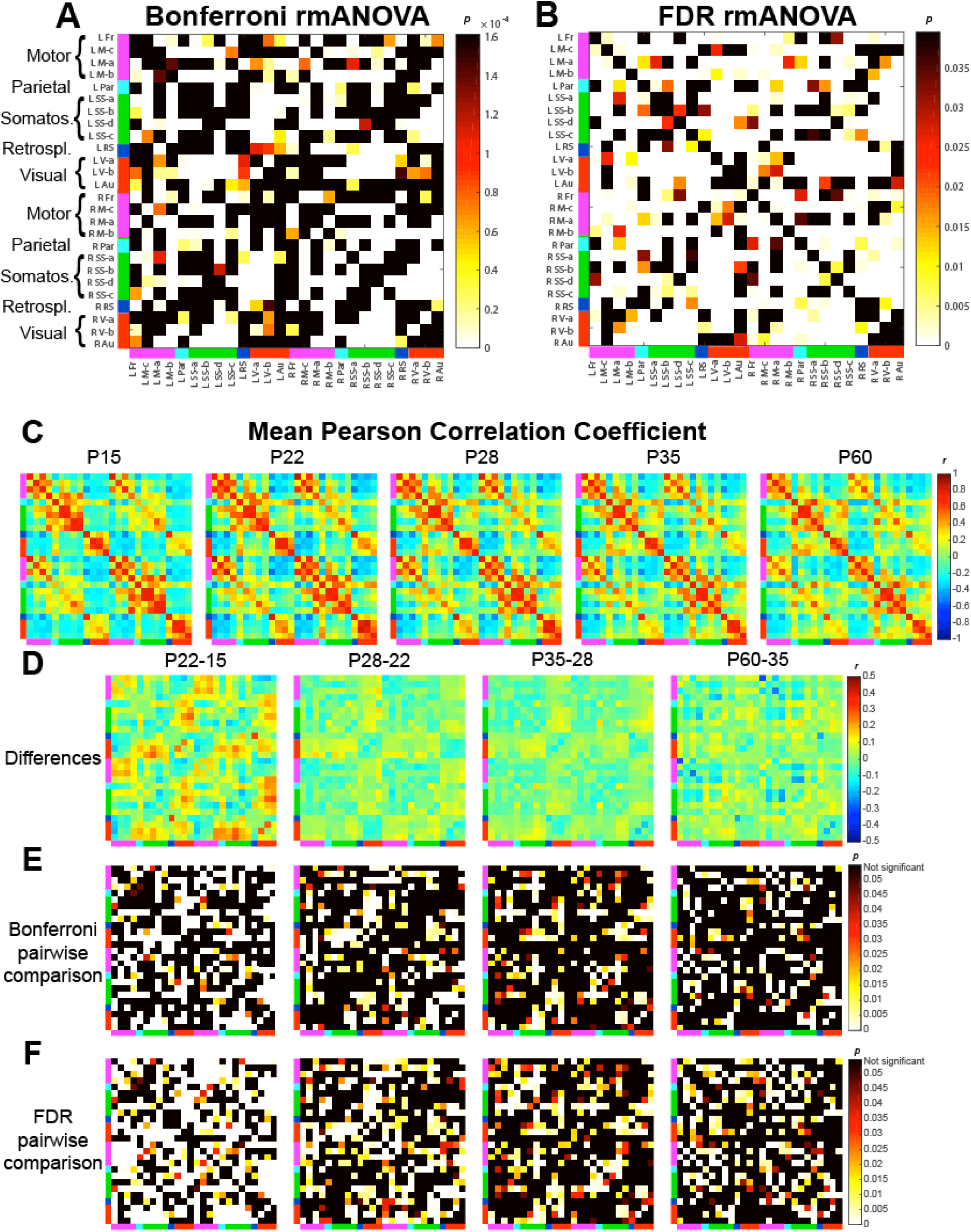
The 325 seed-seed connections between 26 cortical seeds include more than 150 connections that significantly change between P15 and P60 with the strongest change between P15 and P22. A-B) Heat maps representing *p*-values of seed pairings that change significantly across time according to a repeated-measures ANOVA after Bonferroni (A) or false discovery rate (B) corrections. Black represents a connection that was not significantly different (significance at *p*<1.54e-4 for a Bonferroni correction and *p*<0.0376 for a false discovery rate (FDR) correction). Motor, Parietal, Somatos (somatosensory), Retrospl (retrosplenial), and Visual are approximate regional clusters. Seed abbreviations are the following: L left, R right, Fr Frontal, M-c Motor 3rd seed, M-a Motor 1st seed, M-b 3rd seed, Par Parietal, SS-a Somatosensory 1st seed, SS-b Somatosensory 2nd seed, SS-d Somatosensory 4th seed, SS-c Somatosensory 3rd seed, RS Retrosplenial, V-a Visual 1st seed, V-b Visual 2nd seed, Au Auditory. C) Mean Pearson correlation coefficient for each time point from left (P15) to right (P60). D) Mean difference in Pearson correlation coefficient between consecutive time points with the earlier time point subtracted from later. E-F) Post hoc pairwise comparisons between consecutive time points of seed-seed connections that were significant in the rmANOVA represented in A (E) and B (F) (n=17).

### Node degree connectivity increases between P15 and P22

Since FC seeds are traditionally defined based primarily on adult literature, we also examined node degree on a pixel-by-pixel basis to allow a holistic and unbiased approach to defining the pattern of FC development and overall extent of connections. Node degree displays the number of functional connections to other pixels, between 0 and 2422 (number of total pixels in the field of view), for each pixel at each time point, with a functional connection defined as having z(r)>0.3. We hypothesized that there would be a steady increase across time as networks fully formed, with region-specific differences such as an extension of node development later in the frontal cortex, as underlying circuits continue to mature. We found that there was a significant change across time in the majority of cortical regions between P15 and P22, with more localized change between subsequent time points (Figure 4). Anterior regions, including frontal cortex, displayed a significantly different node degree in the P22-P28 and P28-P35 period and between P35 and P60 to a more limited extent. Retrosplenial node degree significantly changed between P35 and P60.

**Figure 4:**
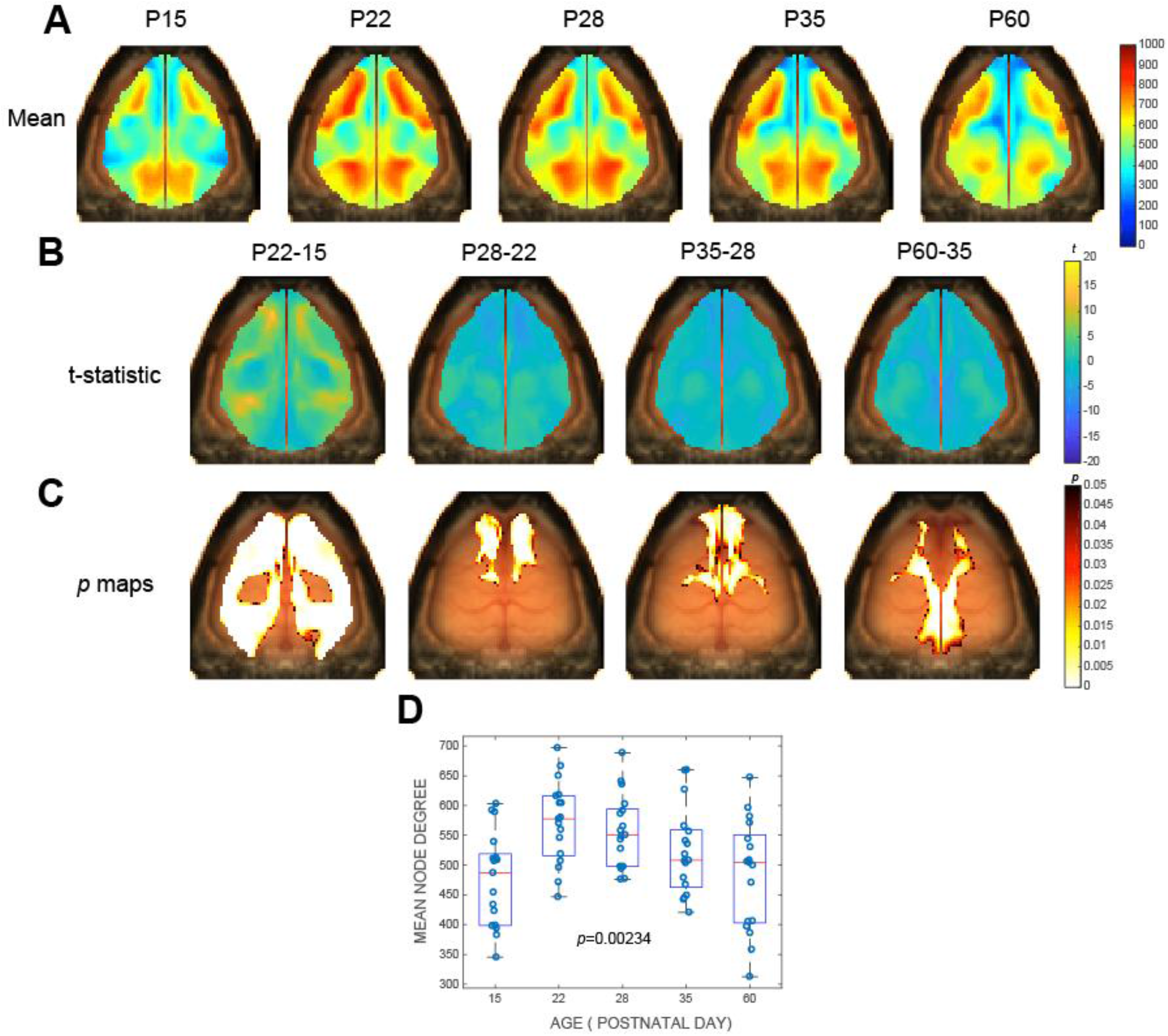
Node degree differs significantly across development with large changes observed between P15 and P22. A) Mean cortical map of the number of other pixels in the field of view with a correlated time course (defined as z(r)>0.3) for each individual pixel. B) t-statistic maps of unpaired t-tests between consecutive time points. C) Pixels with significant homotopic contralateral FC changes between consecutive timepoints following a cluster-based multiple comparison correction. D) Box plots of node degree across time and scatterplot of individuals’ node degree. *p*-value represents rmANOVA result (n=17, 0.4-4.0 Hz).

### Developmental FC change is dependent on anterior-posterior axis location

After examining developmental trajectories of individual seed pairs and pixels, we sought to examine the data set for novel, broad cross-regional patterns of development across the cortex. Some studies have previously reported a temporal progression from posterior to anterior regions in the development of structural connections that are both gray and white matter-based (Gervan et al., 2017). As we observed the peak of most of the connections to occur at P22, we therefore divided seed connections into those positive (correlated) and those negative (anti-correlated) at P22 and explored if there were large-scale trends in how these connections developed. We found that the seeds correlated with each other at P22 tended to display an overall decrease in FC between P15 and P60 in anterior regions and displayed a gross increase in FC in posterior regions (Figure 5). Meanwhile, functional connections anticorrelated at P22 largely increased in strength between P15 and P60 as they became relatively more correlated. Taken together, these changes indicate that there may be a discrete anterior-posterior separation in direction of FC developmental change in functional connections already positively correlated with each other early in development.

**Figure 5:**
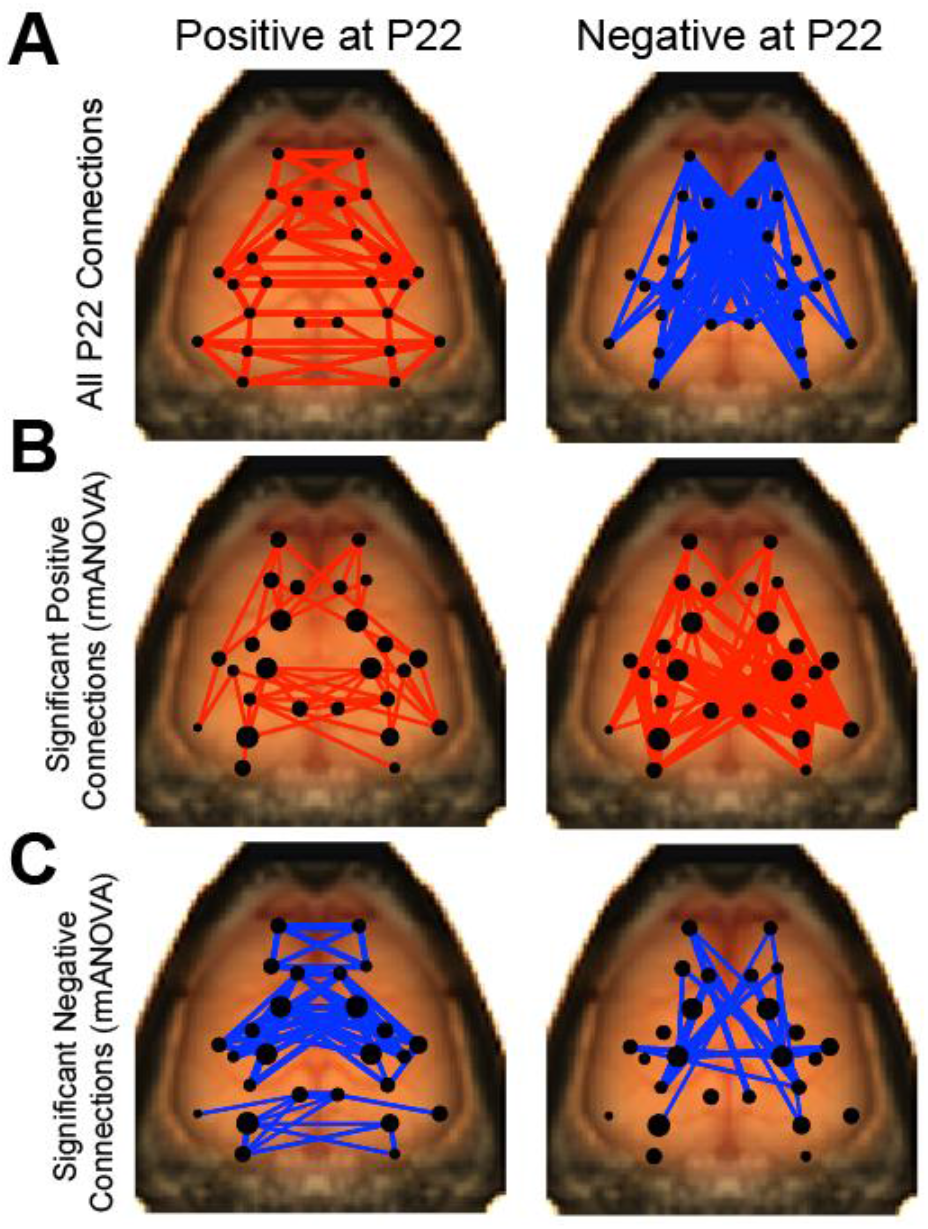
Functional connections significantly changed across development largely showed a decrease in strength if in anterior regions and an increase in posterior regions if positively correlated with each other at P22. **A)** The 50 most correlated (left, red) and 50 most anticorrelated (right, blue) seed-seed connections at P22 and the associated seeds’ positions. **B-C)** All seed-seed connections which were significantly different across time in a rmANOVA with Bonferroni correction (Figure 3) were graphed if the seeds were correlated (left) or anticorrelated (right) at P22 with B) connections that had a net increase in FC and C) connections that had a net decrease in FC plotted separately. Connections that increased in strength between P15 and P60 are represented by red and those that decreased by blue. Size of seeds represents the number of functional connections that seed had which were significant. Thickness of lines represent their significance rank within the multiple comparisons (thicker line=more significant) (n=17).

### Shift in local-distal connection strength occurs between P15 and P22

In order to determine more cortex-wide changes in development, we investigated if the association of distance and FC strength between seeds varied across time. When examining distal/local FC relationships, we hypothesized that based upon a previously documented increase in distal/local FC ratio over time (Ma et al., 2018), we would see a significant increase across development in how highly the more distal connections are represented within a highly-correlated set of connections. We therefore plotted the distance of all seed-seed connections compared to mean Pearson’s correlation coefficient at that time point. We established that there was a clear relationship between distance and FC strength in all seed-seed connections (Figure 6A), but potential differences in that relationship were subtle across age (Figure 6B). We therefore specifically analyzed just the strongest connections to determine if they change on average, by analyzing the mean distance between FC connections above a minimum threshold of correlation (z(r)>0.3)). We found that the mean distance between above-threshold seed-seed connections increased between P15 and P22 but did not significantly change between subsequent timepoints.

**Figure 6:**
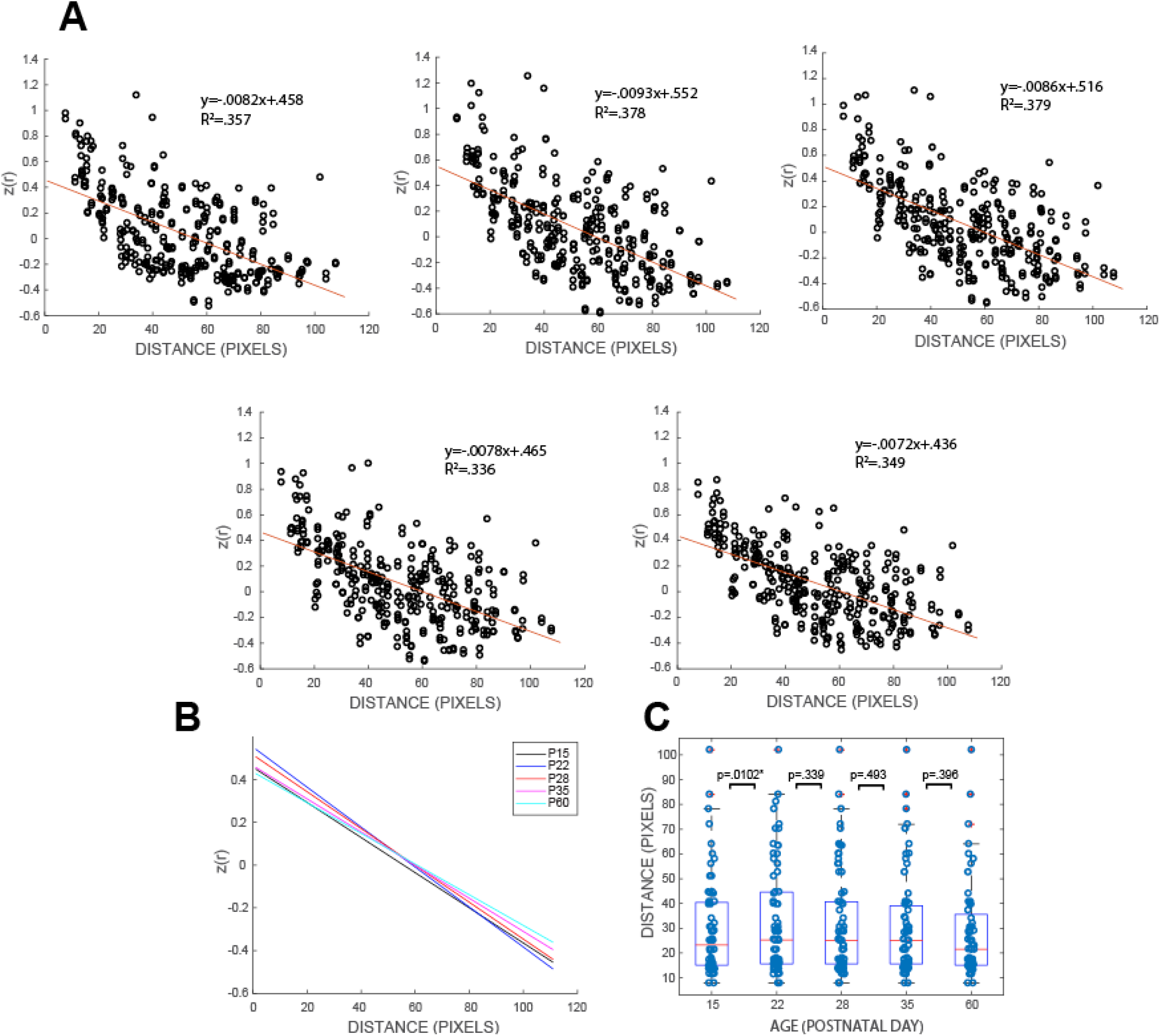
Distance-based distal/local FC change is most evident between P15 and P22. **A)** Scatterplots of mean distance of each seed-seed functional connection within a 26-seed cortical seed set. B) Comparison of slopes between time points. B) Comparison of best-fit lines for the relationship of connections’ z(r) and distance at each developmental age. C) Boxplot of distance between each seed-seed pair with mean FC strength z(r)>0.3 at each time point. *p*-values represent a Wilcoxon rank sum test when comparing distances from each mouse’s connections that pass the z(r)>0.3 threshold (n=17 mice).

### Intrahemispheric connections are more dominant early in development

Finally, as changes to callosal structure are well-documented during this period of development, we sought to determine whether the balance of FC strength between and within hemispheres changed across time. This investigation of hemispheric connection rebalancing also served as a final proof of principle analysis, here addressing whether callosal structural development is paralleled by changes in functional connection dominance. Previous studies have shown an increase in the distal/local FC ratio across development but not examined what a strict differentiation between within- and between-hemisphere connections might reveal. We found that the mean strength of contralateral connections compared to ipsilateral FC increased after P15 (Figure 7), indicating that ipsilateral connections became relatively less dominant when compared to contralateral connections following the third postnatal week. After P22, this contralateral/ipsilateral FC ratio stabilized and did not continue to increase.

**Figure 7:**
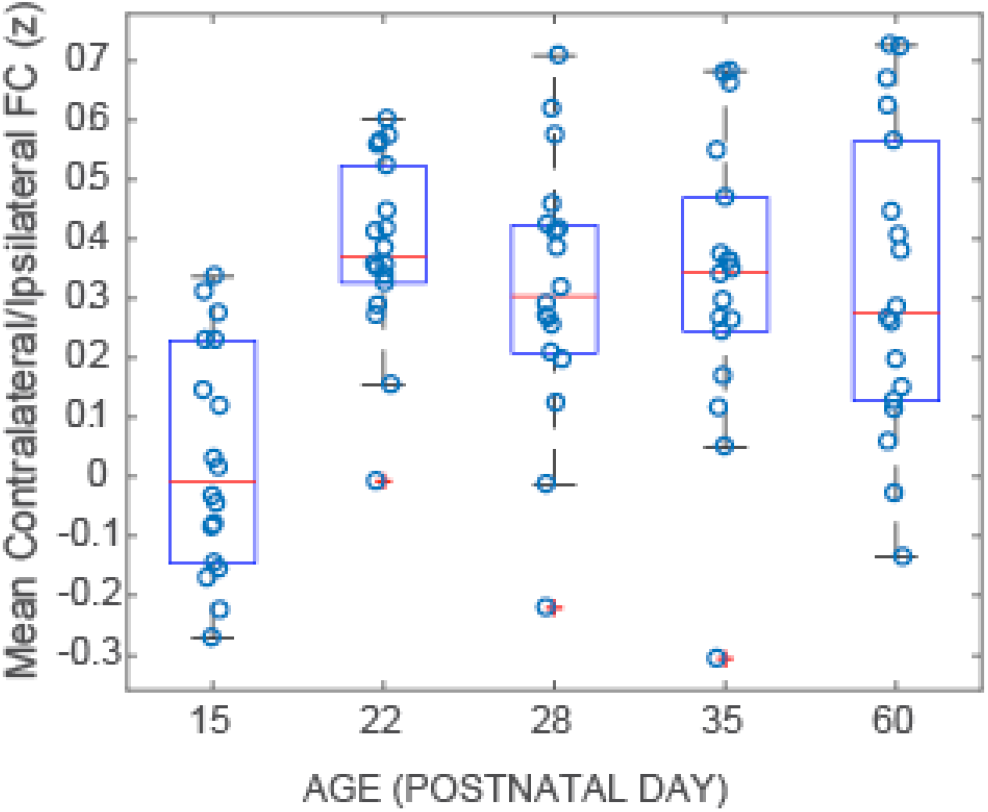
Intra- and interhemispheric FC changes across time during development. Mean contralateral/ipsilateral FC ratio when all 325 connections are designated as within- or between-hemisphere connections (rmANOVA, *p*=1.18e-04, n=17).

## Discussion

Using calcium fluorescence as a direct measure of neural activity, our study has produced a dataset documenting cortical functional connectivity development in the mouse. We have established a FC resource for the research community that provides the opportunity for a wide variety of questions about functional brain development to be asked and answered, and have provided a variety of proof of principle analyses characterizing some aspects of the development of functional connectivity, for example by asking and answering a question regarding how the relationship between intra- and interhemispheric FC develops as structural callosal changes occur. Furthermore, our imaging of the same mice at 5 time points starting at P15 demonstrated that longitudinal and early development imaging is feasible and can shed light on within-individual differences in ways that cross-sectional studies cannot. Finally, calcium imaging allows us to make statements more definitively about neural activity without the potential confound of neurovascular coupling changes.

In our examination of large-scale changes between P15 and early adulthood in the mouse, our study found that various functional connections significantly change across time, and that the developmental trajectories of those connections are highly region-specific. The significant developmental changes we saw in FC allow us to provide a better understanding of the functional correspondences of the well-documented structural changes findings previously reported in the literature. We found that functional and structural connectivity typically closely mirror each other, but FC allowed us to also identify regional coactivations even when direct structural links do not exist (Greicius et al., 2009; Honey et al., 2009; Koch et al., 2002). Our results indicate that function is refined during this period from eye opening to early adulthood, reflecting similar structural changes in the white matter tracts and axonal and synaptic distribution that are reported in the literature. The early rise in FC strength observed between P15 and P22 in a majority of connections may reflect continued synaptogenesis in this early time period, while the later decline in FC strength observed in many regions may be attributable to the pruning of exuberant projections in adolescence. Visual development traditionally proceeds during a time period included in our reported data points. We observe a rise in calcium FC between P15 and P22 following eye opening and then a fall during the visual critical period between P21 and P35. Immediately following eye opening at approximately P15, bilateral visual connectivity increased by approximately 55% by P22, while between P22 and P35, that bilateral visual FC decreased by approximately 60%. This mirrors known changes in the visual cortex during the critical period for ocular dominance plasticity, which extends from P21 and P35 with a peak around P26 and reflects increased GABAergic inhibition observed in layers 2-3 (Dräger, 1975; Espinosa and Stryker, 2012; Kannan et al., 2016; Wagor et al., 1980). The decrease in bilateral connectivity observed may represent pruning of synapses and projections or structural refinements within the visual cortex previously observed in structural development and other functional studies as the critical period progresses and visual function is refined. Competition between eyes in the mouse visual cortex induces dendritic spine refinements in excitatory pyramidal neurons during the critical period (Mataga et al., 2004). Our GECI is expressed in excitatory pyramidal neurons in layers 2-3, thus our findings of altered calcium FC may reflect specifically this excitatory refinement. Kraft et al. also reported developmental FC change dependent on eye suturing that suggested the P22-P35 period is critical to visual development (Kraft et al., 2017). By utilizing GECIs rather than hemodynamics, and expanding to additional time points, we have therefore expanded upon structural and functional knowledge of the visual critical period and directly traced the development of unperturbed visual FC throughout this time of dynamic change.

We also identified groups of unexpectedly similar trajectories between certain cortical regions’ homotopic connectivity, where developing regions acted distinctly yet shared some similarities. Somatosensory regions showed little dynamic change in bilateral FC across time, which may suggest that somatosensory connections are already well-established before P15, as infant mice extensively use that circuit. Frontal regions’ decline in connectivity could represent the later pruning that has previously been observed in the prefrontal cortex in numerous studies (Huttenlocher & Dabholkar, 1997; Mallya et al., 2019). The relatively slow pacing of myelination, which is known to occur into adulthood, may also exert a strong influence on FC and explain why some regions unexpectedly share or lack similar developmental trajectories despite expectations based on the region’s functional similarity or dissimilarity with each other.

The extreme changes between P15 and P22 over the entire cortex, as well as anterior-specific regional change at later time points, is especially evident when pixel-wise analysis of bilateral functional connectivity and node degree was performed. By expanding beyond just a canonical seed set, we were able to identify pixel groups that survived a cluster-based correction and therefore unbiasedly identified other changing regions in the developing brain. The strong increase in both bilateral FC and node degree in almost the entire dorsal cortex suggested that this earliest change reflects a brainwide phenomenon where the third postnatal week sees a discrete and enduring change to a state much more similar to the adult functional connectome. The small portions of the functional connections we sampled that did not significantly change in this earliest time period interestingly include the homotopic contralateral somatosensory connections, which link functional regions that are known for having a critical period earlier than P15, the earliest time point we sampled (Reh et al., 2020). Following P22, consistent changes in bilateral FC and node degree were seen in the anterior portions of the brain that include frontal cortex, an extended developmental change which aligns with the fact that frontal and specifically prefrontal cortices are well-known for their later maturation (Miller et al., 2012; Snaidero and Simons, 2014). Besides the frontal cortex, the motor, visual, and retrosplenial cortices significantly changed between P22 and P60 when the bilateral pixel maps were examined. Meanwhile, following a large increase in node degree across the majority of the brain between P15 and P22, anterior regions had significant node degree alterations between P22 and P60, while the retrosplenial region’s node degree differed only between P35 and P60 after the original P15-P22 period of greatest change. While continued change in the frontal cortex would be expected for the previously discussed reasons, the fact that motor retrosplenial regions display post-P22 differences could reflect some of the refining of connections previously reported in studies documenting synaptic pruning occurring later in association cortex than sensorimotor cortex (Huttenlocher and Dabholkar, 1997; Levitt, 2003). Motor FC’s change after the P15-P22 period is perhaps most puzzling because motor cortex is traditionally thought to be among the earliest to develop; our findings may reflect an increased refinement of FC as complex motor skills develop and complex behaviors are learned that require motor coordination with other senses or processes.

Finally, a rebalancing of global connections and dominant characteristics is also reflected in the progression of FC development over the period we examined. Homotopic contralateral FC and local connection dominance both tended to decrease across time, suggesting that more complex relationships between more distinct or distant regions tend to become more dominant with age. The change across time in seed-seed correlation strength also appeared to be partially segregated spatially into connections that gained FC strength (posterior) and those whose correlation coefficients declined between P15 and P60 (anterior), which does not exactly align with the posterior-to-anterior progression of myelination in the brain but nonetheless brings up the possibility that the anterior-posterior axis plays an important role in functional development. On the whole, there was a strengthening of strong connections, as the overall local-distal ratio of connections shifted between P15 and P22 to have more distal connections within the set of functional connections that exhibited above-threshold strength. We also observed that there was a shift in inter- and intrahemispheric connection strength between P15 and P60 as intrahemispheric connections became less dominant. These results expanded upon what was previously reported regarding distal/local connection ratios increasing with age (Ma et al., 2018). Overall, this suggests that there is a gross shift from a region’s close connections with functionally or geographically similar neighbors and toward complex relationships with other regions with more distant functions or locations, and that this shift may be specifically linked to the period between P15 and P22.

This functional connectivity dataset therefore will provide a highly useful resource to the field, to answer questions that are both natural extensions of this work as well as more novel applications. This study overall supports the conclusion that developmental trajectories of functional connections have region-specific elements as well as some unifying features that could be attributed to a system-wide shift toward more complex, refined regional connections. These findings lay the groundwork for further study of healthy development and define several questions that can be answered using this resource: Namely, whether an excitation-inhibition rebalance might be driving the extreme FC change during the third postnatal week; also, whether node degree and bilateral FC change in motor cortex post-P22 reflects behavioral development of complex motor skills; and additionally, whether there is an anterior-posterior divide in the trajectory of functional regions’ healthy brain development and if so, if it persists in post-P60 development. Additionally, questions regarding how individual mice deviate from the norm during FC development, how neurovascular coupling develops, and how temporal FC patterns such as those identified by lag analysis evolve over this time period. This technique and these findings are also applicable to a wide variety of disease models and are particularly promising for the study of neurodevelopmental disorders, as they facilitate the identification of trajectory abnormalities and characterization of how they diverge from baseline. This dataset also can serve as a resource to the field in the future, to permit temporal and spatial targeting of research questions to ensure maximal impact of cross-sectional studies. This minimally invasive method and the knowledge gained regarding the periods of most dynamic change in each region provide an important resource and springboard to targeted studies of disease in the future.

## Acknowledgements

This work was supported by the National Institute of Neurological Disorders and Stroke (F31NS110222, R.M.R, R01NS099429, J.P.C. and R01NS090874, J.P.C.), National Institute of Mental Health (R01MH124808 and R01MH107515, J.D.D.) and National Institute on Aging (F30AG061932, L.M.B.).

## Author Contributions

Conceptualization, R.M.R., J.D.D. and J.P.C.; Methodology, R.M.R., J.D.D. and J.P.C.; Software, L.M.B., Formal Analysis, R.M.R.; Investigation, R.M.R. and A.R.B.; Writing - Original Draft, R.M.R., J.D.D. and J.P.C.; Writing – Review & Editing, R.M.R., L.M.B., A.R.B., M.D.R., J.D.D. and J.P.C.; Funding Acquisition, R.M.R., J.D.D. and J.P.C.; Resources, L.M.B. and M.D.R.; Supervision, J.D.D. and J.P.C.

## Methods

### RESOURCE AVAILABILITY

#### Lead contact

Further information and requests for resources should be directed to and will be fulfilled by the lead contacts, Joseph Culver and Joseph Dougherty.

#### Materials availability

This study did not generate new unique reagents. Pipeline code is available at (Brier and Culver, 2021). Dataset is available upon request.

### EXPERIMENTAL MODEL AND SUBJECT DETAILS

#### Animals

Mice were housed and experiments were conducted in accordance with the relevant guidelines of the Washington University Institutional Animal Care and Use Committee and the approved Animal Studies Protocol. All mice were maintained on a 12:12h light/dark cycle and provided with freely available food and water. *Thy1*-GCaMP6f (JAX:024276) homozygous males were bred to C57BL/6J (JAX:000664) females to produce *Thy1*-GCaMP6f hemizygous experimental animals, and litters culled at postnatal day 5 to 4 pups. All surviving mice at P15 were imaged. Seventeen mice total (12 males, 5 females) were used for data analysis, with one mouse discarded prior to analysis due to failure to pass data quality thresholds.

### METHOD DETAILS

#### Surgical windowing

A transparent chronic optical window made of Plexiglas was fitted on each mouse at P14 to enable repeated head-fixed imaging during wakefulness. The head was shaved and an incision was made along the midline of the scalp to retract the skin and expose an approximately 1.1 cm^2^ dorsal cortical field of view. The optical window was adhered to the dorsal cranium using Metabond clear dental cement (C&B-Metabond, Parkell Inc., Edgewood, NY) while under isoflurane anesthesia (3% induction, 1% maintenance, 0.5 L/min) and topical lidocaine analgesia. Pups were returned to their mother to recover for approximately 24 hours before imaging data were collected.

#### Fluorescence and Optical Intrinsic Signal (OIS) Imaging

Mice were imaged during their 6 a.m.-6 p.m. light cycle. Mice were headfixed and imaged longitudinally during wakefulness in the calcium fluorescence imaging system at 5 developmental timepoints (P15, P22, P28, P35, and P60). At the earliest time point (P15), a water heater (Gaymar heating/cooling T/Pump system, Braintree Scientific, Inc.; Braintree, MA) was used to help maintain body temperature at 37°C before weaning. Sequential illumination was provided by 4 LEDs (470 nm (GCaMPf excitation), 530 nm, 590 nm, and 625 nm; Mightex Systems; Pleasanton, CA) that passed through dichroic lenses (Semrock; Rochester, NY) into a liquid light guide (Mightex Systems; Pleasanton, CA). A 75 mm f/1.8 lens (Navitar; Rochester, NY) then focused the light from the liquid light guide onto the cortical surface. Video was collected using a sCMOS camera (Zyla 5.5, Andor Technologies; Belfast, Northern Ireland, United Kingdom) directly above the dorsal surface of the head, and fluorescence and reflectance frames were collected at a rate of 16.8 Hz for each of the 4 LED wavelengths, with a 515nm longpass filter (Semrock; Rochester, NY) utilized to filter out GCaMP6f excitation light. Specular reflection artifacts were discarded by use of cross polarization (Adorama; New York, NY) between the illumination and collection lenses. Six 5-minute runs were collected for each mouse in each session. Imaging data were binned to 156×156 pixels^2^ images with each pixel approximately 100 µm^2^.

### QUANTIFICATION AND STATISTICAL ANALYSIS

All analyses were performed in MATLAB 9.3 (R2017b). Data were processed using the previously published optical calcium imaging pipeline (Brier and Culver, 2021) and spatially downsampled to 78×78 pixels^2^ and temporally downsampled by 2 for analysis. Individual 5-minute runs were discarded prior to analysis if raw light levels had a variance greater than 1.5% across the run. Frames within each run were removed if they exceeded a threshold of 1 standard deviation in the global variance of the temporal derivatives (GVTD) in the oxyhemoglobin signal in the 0.4-4.0 Hz frequency band, adapted from the GVTD method of Sherafati et al. (Sherafati et al., 2020). Masking of non-brain pixels was performed according to the previously published methods, with a modification to allow pixels represented in 61 or more of the 85 imaging sessions to be included only for those mice whose field of view encompassed them. All analyses were performed on data filtered to the delta (0.4-4.0 Hz) frequency band (Supplementary Figure 1) and used a cortical seed set based upon and expanded from White et al. (White et al., 2011) (Supplementary Figure 2). Repeated-measures ANOVA with a Greenhouse-Geisser sphericity correction were performed on all analyses examining change across the five timepoints, and paired t-tests on all pairwise comparisons between two timepoints unless indicated otherwise. When performing post hoc comparisons on functional connections to produce Figure 3E-F, seed-seed pairs were only compared that demonstrated significance in the rmANOVA as seen in Figure 3A-B. Otherwise, a false discovery rate (FDR) or Bonferroni correction was performed for multiple comparisons of seed-seed connections, and all figures and analyses examining only the set of significant rmANOVA connections was based upon the Bonferroni-corrected significance threshold. Multiple comparison correction for pairwise pixel-by-pixel bilateral and node degree analyses was performed by dividing the alpha level of 0.05 by the number of pairwise comparisons (four) and then performing a cluster-based correction previously described here (Brier and Culver, 2021). Briefly, a random field theory approach used broadly in the fMRI (Friston et al., 1994) and high-density diffuse optical tomography (HD-DOT) literature (Hassanpour et al., 2014), was applied to handle the multiple comparisons problem and create a cluster-size (units in pixels) threshold to delineate significantly altered cortical regions in a pixel-wise statistical comparison. This approach leverages the spatial dependence of neighboring pixels, either through biological function or imaging system blurring (as quantified through the full-width half-maximum of the spatial autocorrelation of an image), and weights larger clusters to have more statistical significance than smaller clusters with the same peak t-value.

Node degree was calculated by correlating each pixel’s time course with the time course of all other pixels and thresholding at a correlation of z(r)>0.3 for each individual pixel (Hakon et al., 2018). Group maps were calculated by summing inter- and intrahemispheric connections surviving threshold (i.e., node degree). Local-distal connection strength data were represented in Figure 6 by creating scatter plots and using polyfit to graph a best-fit line. Local-distal connection strength was quantified by comparing Pearson z(r) and distance, and unpaired t-tests between consecutive time points were performed by comparing datasets from each time point consisting of the pixel distance between seed centers of each connection with z(r)>0.3 for each individual mouse. Calculations of inter- and intrahemispheric FC changes across time were performed by calculating mean Pearson’s z(r) for each seed’s contralateral connections, averaging this across all seeds to calculate a single contralateral FC value, then dividing it by a single ipsilateral FC value for that same mouse, derived from all seeds’ ipsilateral (intrahemispheric) connections. Each mouse’s contralateral/ipsilateral FC ratio was then compared across time in a rmANOVA.

## Supplemental Information

**Supplementary Figure 1:**
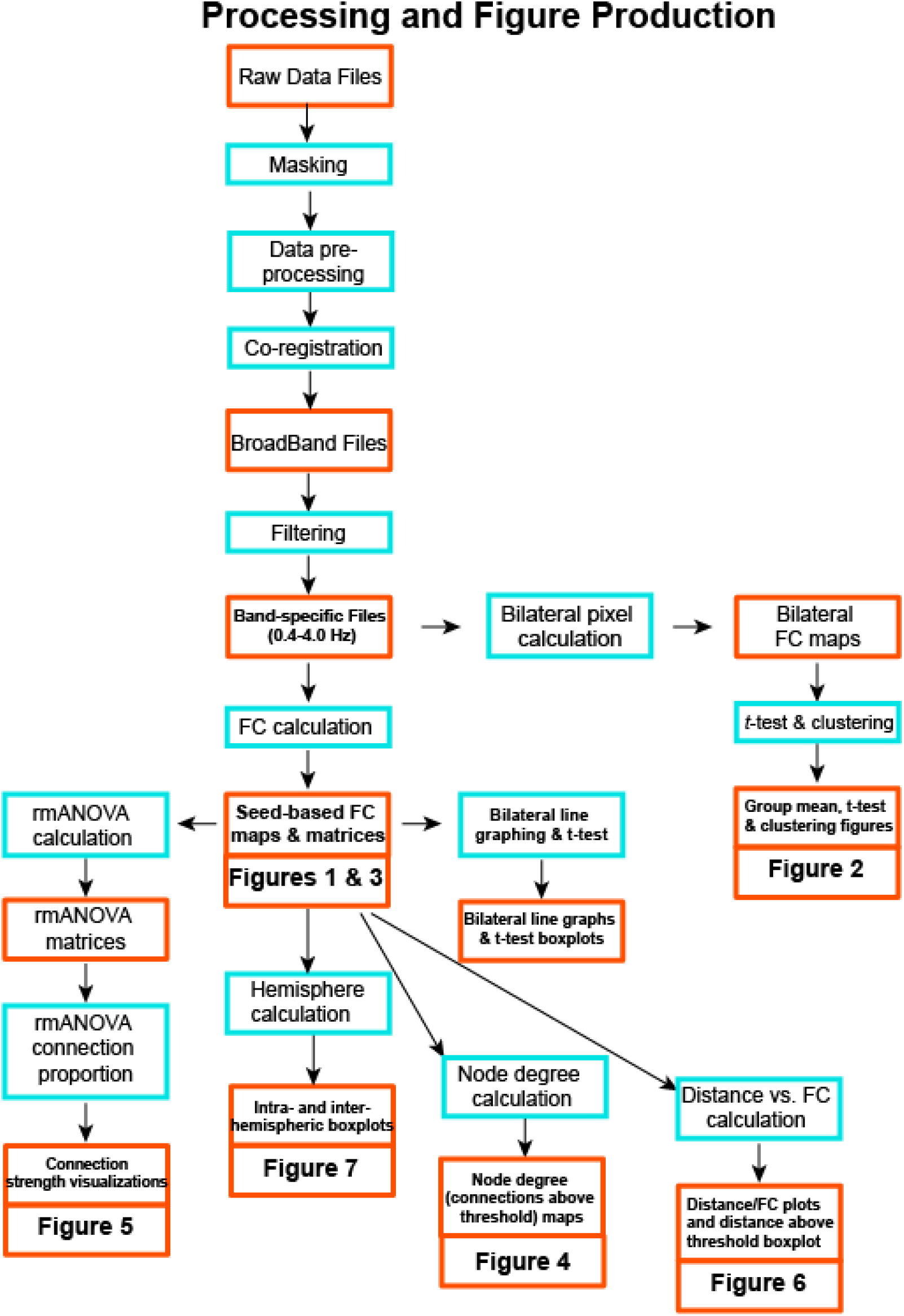
Data processing for calcium fluorescence data and functional connectivity analyses. Blue boxes indicate computational steps, while red boxes represent resultant files or figures. All processing scripts available at (Brier and Culver, 2021) and on request.

**Supplementary Figure 2:**
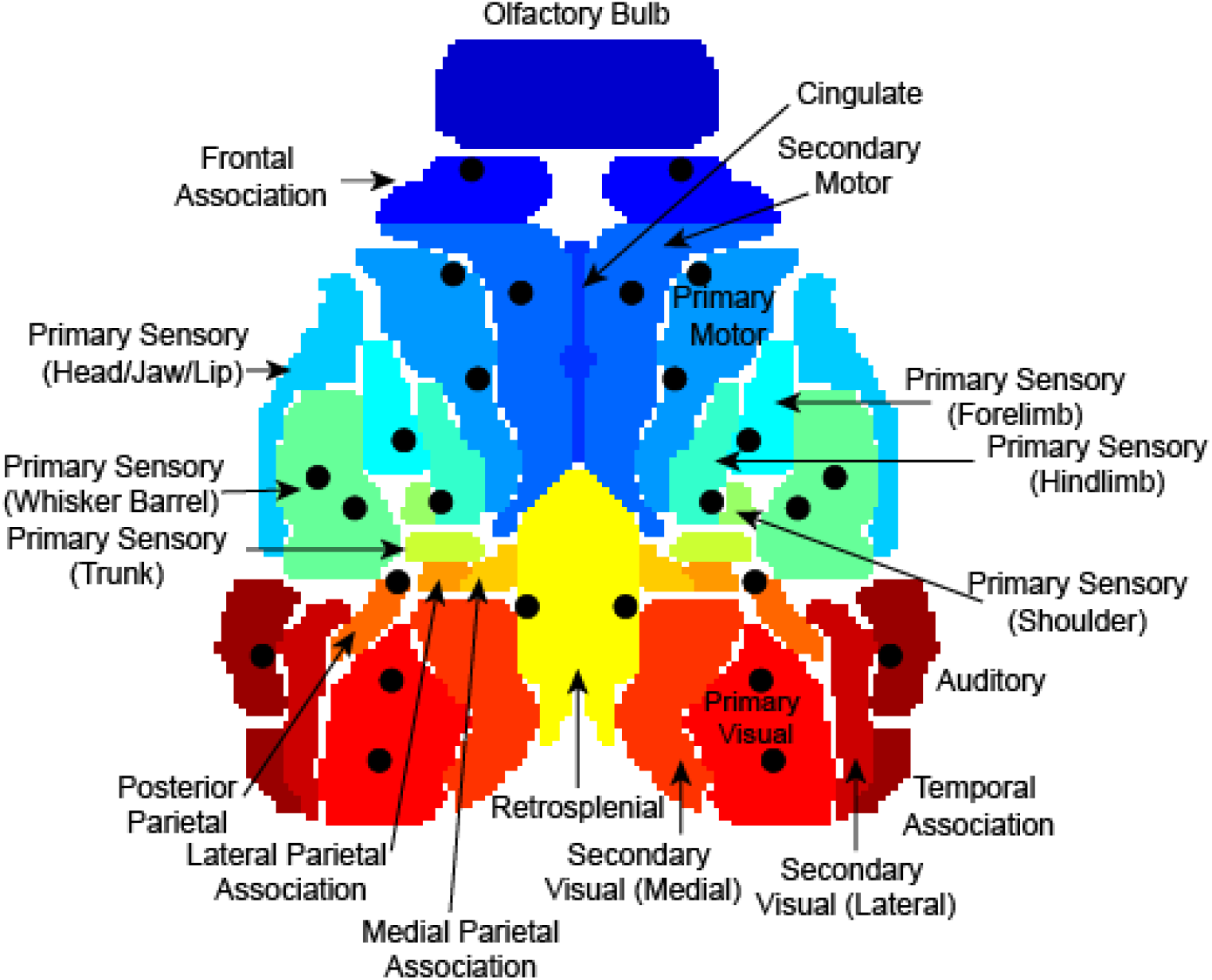
Paxinos atlas-based map of functional regions and sampled seeds adapted from White et al. (2011). All regions are identical between the hemispheres when reflected across midline.

